# MetaFun: Unveiling Sex-based Differences in Multiple Transcriptomic Studies through Comprehensive Functional Meta-analysis

**DOI:** 10.1101/2021.07.13.451905

**Authors:** Pablo Malmierca-Merlo, Rubén Sánchez-Garcia, Rubén Grillo-Risco, Irene Pérez-Díez, José F. Català-Senent, Santiago Leon, Helena Gómez-Martínez, María de la Iglesia-Vayá, Marta R. Hidalgo, Francisco Garcia-Garcia

## Abstract

While sex-based differences in various health scenarios have been thoroughly acknowledged in the literature, we lack a deep analysis of sex as a variable in this context. To fill this knowledge gap, we created MetaFun as an easy-to-use web-based tool to meta-analyze multiple transcriptomic datasets with a sex-based perspective to gain major statistical power and biological soundness. Furthermore, MetaFun can be used to perform case-control meta-analyses, allowing researchers with basic programming skills to access this methodology.

**Availability and implementation:** MetaFun is freely available at http://bioinfo.cipf.es/metafun The back end was implemented in R and Java, and the front end was developed using Angular

**Supplementary information:** R code available at https://gitlab.com/ubb-cipf/metafunr

## 1 Introduction

Sex-based differences in different health scenarios have been thoroughly acknowledged in the literature [1,2]; however, this variable remains incompletely analyzed in many cases. Studies often neglect sex as a variable when considering the experimental design of studies, leading to experiments with samples of just one sex in extreme cases. As a result, the underlying mechanisms behind sex-based differences in many diseases and disorders remain incompletely established.

Fortunately, the scientific community has worked to significantly improve this situation in recent times, and researchers have begun to include the sex perspective in their research; however, a vast amount of generated data currently stored in public databases (such as Gene Expression Omnibus (GEO) [3] or NCI’s Genomic Data Commons (GDC) [4]) remains unanalyzed from this perspective. The information in these databases represents a powerful resource that must be considered.

When exploiting these resources with a particular objective, multiple studies dealing with similar scientific questions can provide different and often contradictory results. No one study is likely to provide a definitive answer; therefore, integrating all datasets into a single analysis may provide the means to understand the results. Defined for this purpose, meta-analysis is a statistical methodology that considers the relative importance of multiple studies upon combining them into a single integrated analysis and extracts results based on the entirety of the evidence/samples [5,6,7]. Unfortunately, applying advanced statistical techniques such as meta-analysis often remains out of reach for biomedical researchers aiming to analyze their data in a straightforward manner.

We designed the “MetaFun” tool to simplify the analytical process and facilitate the application of functional meta-analysis to researchers working with multiple transcriptomic datasets. Meta-analysis approaches can analyze datasets from perspectives such as sex and combine datasets to gain significant statistical power and soundness. MetaFun is a complete suite that allows the analysis of transcriptomics data and the exploration of the results at all levels, performing single-dataset exploratory analysis, differential gene expression, gene set functional enrichment, and finally, combining results in a functional meta-analysis.

## 2 Methods

The MetaFun tool is available at https://bioinfo.cipf.es/metafun while additional help can be found at https://gitlab.com/ubb-cipf/metafunweb/-/wikis/Summary.

### 2.1 Input Data and Experimental Design

MetaFun takes a set of at least two CSV expression files and two TSV experimental design files as inputs. CSV expression files must include normalized transcriptomics data from comparable studies with assimilable experimental groups. Columns must contain the study samples, while rows must contain analyzed genes as their Entrez Gene ID. The first row contains sample names. TSV experimental files define the class to which each sample of the study belongs by including at least two columns: sample names and the class to which they belong. Column names in the CSV expression files must match row names in the corresponding TSV experimental file. Accepted reference organisms are (for the moment) humans (*Homo sapiens*), mice (*Mus musculus*), and rats (*Rattus norvegicus*). Analyses can be made with respect to a comparison that must apply to all datasets. Options include the classical comparison - *Case* vs. *Control* (**Fig. 1A**) - or a sex-perspective comparison - (*Male case* vs. *Male control*) vs. (*Female case* vs. *Female control*) (**Fig. 1B**) - in which the effect under study is compared between sexes.

**Figure 1:**
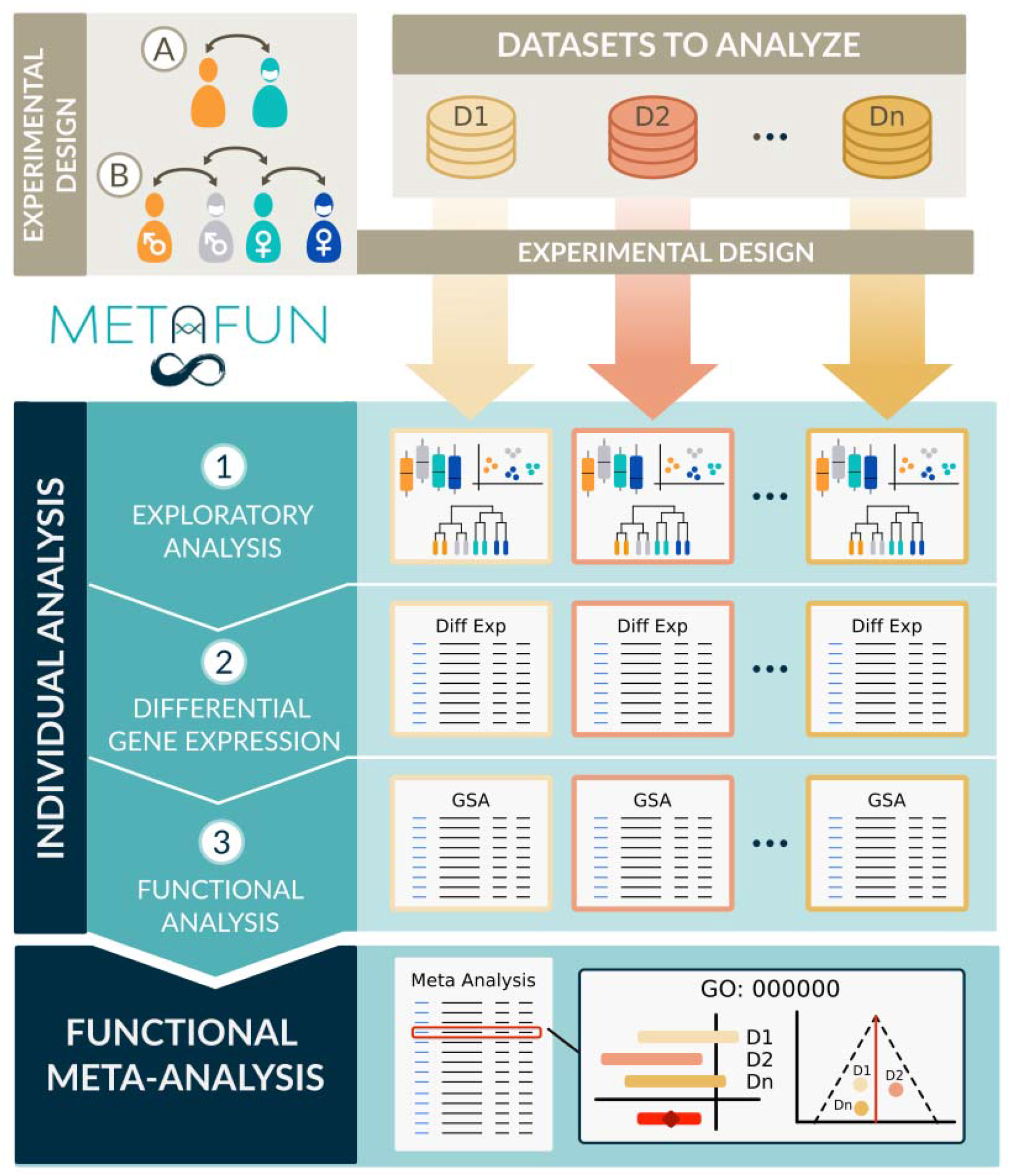
MetaFun pipeline. First, datasets and experimental designs are uploaded as CSV and TSV files. Available comparisons include (**A**) a classical comparison - *Case vs. Control* - and (**B**) a sex-perspective comparison - (*Female case* vs. *Female control*) vs. (*Male case* vs. *Male control*). Single experiment analyses include 1) exploratory analysis, 2) differential gene expression, and 3) functional analysis performed on each dataset. Finally, the results are integrated into a functional meta-analysis. The MetaFun tool allows users to explore all results generated during the process.

### 2.2 Single Dataset Analyses

After the selection of the studies and experimental design, MetaFun analyzes each dataset separately with an individual analysis consisting of:

□ an exploratory analysis including boxplots, PCA, and cluster plots using the *plotly* library [8]

□ a differential gene expression analysis using the *limma* package [9]

□ a gene set enrichment analysis (GSEA) [10] based on gene ontology (GO) [11] from the *mdgsa* package [12].

Figures and tables from these analyses can be explored and downloaded from the *Results* area once the job has been completed. Links to NCBI [13] and QuickGO [14] databases are present to detail the results.

### 2.3 Functional Meta-analysis

MetaFun combines the gene set functional enrichments from all datasets into a meta-analysis with the same experimental design using the *metafor* package [15]. Forest and funnel plots are generated utilizing the *plot*.*lyJS* library [8]. Figures and tables from this meta-analysis are interactive and may be explored and downloaded from the *Results* area once the job has been completed.

### 2.4 Implementation

The MetaFun back end was written using Java and R and is supported by the non-relational database *MongoDB* [16], which stores the files, users, and job information. The front end was developed using the *Angular* framework [17]. All graphics generated in this web tool were implemented with *Plot*.*ly* [8] except for the exploratory analysis cluster plot, which uses the *ggplot2* R package [18].

### 2.5 Study Cases

As an example, MetaFun includes two sets of pre-selected study cases, one for each accepted species: human and rat. The study cases can be executed directly from the web tool, allowing the tool’s functionalities to be easily explored. The human study case includes nine studies from lung cancer patients [6].

## Acknowledgments

The authors thank the Principe Felipe Research Center (CIPF) for providing access to the cluster, which is co-funded by European Regional Development Funds (FEDER) in Valencian Community 2014–2020. The authors also thank Stuart P. Atkinson for reviewing the manuscript and Jaime Llera, Fernando Gordillo, Violeta Lillo, Aaron Garcia, Miriam Ferreiro, Miguel Ferriz, and Daniel Gonzalez for evaluating the web tool.

## Supplementary Material

### S1. Web Tool Overview

The web tool can be used with registered or anonymous users. Registered users will keep their data and jobs stored from one session to the other, while data and jobs from anonymous users will not be saved after leaving the session. The general design of the web tool includes an upper right menu with basic tool functionalities, a left side panel with specific submenus, and a central panel from which to interact with the web. Users are directed to a form launching a new job after logging in, which can be otherwise accessed through the *New Analysis* button in the top right menu. The *New Analysis* form passes through a series of steps, asking for information that must be completed (see **Supplementary Section S2** for details) and allows for a new meta-analysis. After the launch and execution, the job will be listed in the jobs area, which can be accessed through the *My jobs* button located in the top right menu. All created jobs are listed and can be accessed through the left side submenu to visualize results. Users can access their personal area through the top right panel using a button carrying their username. Each user’s personal area includes a browser for folders and information regarding all launched jobs. The user’s personal area submenu supports a series of actions related to personal settings and deleting options. The top right panel also includes an exit icon button that logs users out and a question mark icon that opens documentation pertinent to the web tool (accessible through https://gitlab.com/ubb-cipf/metafunweb/-/wikis/Summary).

### S2. Input Data

All datasets in the same meta-analysis should be comparable, including similar experimental designs and individuals with similar conditions. At least two datasets must be included in a meta-analysis. Input data consists of one expression matrix and one experimental design file for each dataset in the meta-analysis. The expression matrix must have been normalized, with samples in columns and Entrez ID genes in rows. The experimental design file must indicate the original group to which each sample belongs, with samples in rows and groupings in columns. More than one grouping per file is accepted, placing each grouping in a different column (for instance, the first column could be sex, the second column experimental conditions, and the third column disease grade). However, only one grouping at a time will be used in a meta-analysis.

### S3. Launching a Meta-analysis

The *New Analysis* button in the top right menu directs users to a form that launches a new meta-analysis. The first tab of the form - labeled *Files* - includes a browser for user files and allows users to upload and manage the datasets to analyze. The *Options* tab allows users to specify the *Effect Model* to *random* or *fixed*, to select the reference organism among *Homo sapiens*, and *Rattus norvegicus*, to define the GO ontologies to analyze (*Biological Process, Molecular Function*, and *Cellular Component*), and whether to propagate the annotation. A brief description accompanies each option to help users in their decisions. The *Studies* tab is used to select the studies to meta-analyze and the experimental design. Depending on the case, selections are made by dragging files from the right panel entitled *My Files* to the columns *Expression* or *Experimental Design*. Matched studies and experimental designs must be placed in the same row, verifying their compatibility by checking sample names in the expression and experimental design files. Users can specify the comparison to perform in the *Comparison* tab. Two different options are currently available: the classical *Case vs. Control*, which compares the effect of a variable, and a sex-based comparison - *(Case Female vs. Control Female) vs. (Case Male vs. Control Male)* - which compares the effect of a variable in females with respect to the effect in males. In the second case, significant results refer to differential effects between males and females and may not coincide with results from the first comparison. After selecting the comparison, users must indicate which study samples are included in each canonical compared group (*Case, Control, Case Female*, etc.) by assigning one of the classes in the experimental design of each study to these canonical groups. Finally, the *Launch* tab contains a summary of the defined meta-analysis, which may be launched through the *Launch job* button after the name assignment.

### S4. Analysis Summary

After the execution of the job and its selection in the left side panel of the *My Jobs* panel, the

*Analysis summary* tab will show a summary of the main results, which include:

□ selected analysis options - name, comparison, effect model, functional profile, and reference organism

□ a table and an interactive barplot describing the number of samples per dataset and per group

□ a table describing the number of differentially expressed genes in each dataset - per column, the studies, total number of analyzed genes, total number of significantly affected genes, number of significantly upregulated genes, and number of significantly downregulated genes

□ a table including the same columns describing the number of significant functional profile items in each dataset - either enriched functions or differentially activated subpathways, depending on the selected functional profile

□ a table including the same columns describing the number of significant functional terms in each ontology - BP for Biological Process, MF for Molecular Functional, and CC for Cellular Component - of the meta-analysis

### S5. Exploratory Analysis

The *Exploratory analysis* tab contains the figures from the unsupervised exploratory analysis performed on each dataset in the meta-analysis. This analysis includes a boxplot representation of the expression of the samples, a clustering of the samples, and a principal components analysis (PCA) plot representing the first two components of the PCA. All samples are colored by the experimental design selected in the meta-analysis.

### S6. Differential Expression

The differential expression analysis is performed with the *limma* library [9], applying *lmFit, contrast*.*fit*, and *eBayes* functions while considering whether samples are paired or unpaired. Results will be displayed as a table in the *Differential expression* tab of the job. The table shows the Entrez ID, Gene Name, the logarithm of the fold-change (logFC), test statistic, raw p-value, and Bonferroni-Holm [19] adjusted p-value of each analyzed feature. The raw p-value initially orders the table, but buttons on column names allow users to order the table differently. Links from the Entrez ID column direct to the NCBI gene database of the specific gene. Different tools allow users to search, download, and filter the table by a maximum p-value.

### S7. Gene Set Enrichment Analysis

The functional analysis consists of a GSEA [10] based on the BP, MF, and CC ontologies from GO [11] defined by users. The pipeline, performed with the *mdgsa* library [12], splits the ontologies, propagates the annotation (if indicated), filters too generic (more than 500 annotated genes) or too specific (less than ten annotated genes) annotations, transforms the p-value into an index, and performs the corresponding comparisons. Results will be displayed as a table in the *GSA* tab of the job. Three subtabs on the top right of the table separately show the results for the three different ontologies. For each ontology, the table shows the GO ID, GO term, the logarithm of the odds-ratio (LOR), raw p-value, Bonferroni-Holm adjusted p-value, and the number of genes included in each analyzed feature. The raw p-value initially orders the table, and buttons on column names allow users to order the table differently. Links from the GO ID column direct to the QuickGO [14] entry of the specific term. Different tools allow users to search, download, and filter the table by a maximum p-value.

### S8. Meta-analysis

The functional meta-analysis integrates the functional analysis results and is performed using the *rma* function of the *metafor* package [15]. For each function, a meta-analysis combines the level of overrepresentation (LOR) of that function in different studies. Two methods have been implemented to perform meta-analyses: the fixed effects models (FE) and the random effects models (DL DerSimonian & Laird; HS Schmidt & Hunter; Hedges, HE) [15]. The fixed effect model has been designed for similar studies (i.e., with the same technology, platform, and at similar times), while the random effect model allows for more significant variability. Results will be displayed as a table in the *Meta-analysis* tab of the job. The table shows the GO ID, GO term, LOR, confidence interval of the LOR, raw p-value, and Bonferroni-Holm [19] adjusted p-value of each analyzed feature. The raw p-value initially orders the table, and buttons on column names allow users to order the table differently. Links from the GO ID column direct to the QuickGO [14] entry of the specific term. Different tools allow users to search, download, and filter the table by a maximum p-value.

### S9. Study Case

The following case describes the potential use of MetaFun in characterizing sex-based differences in lung adenocarcinoma. The results obtained were published in [20].

#### Input data

Each study requires two files: one with expression data and a second with the description of the experimental groups to which each sample belongs, indicating the sex of the participant. The files corresponding to this use case can be downloaded at the following link - https://gitlab.com/ubb-cipf/metafunpipeline/-/blob/master/Homo%20Sapiens%20(Adenocarcinoma).zip

#### Four Simple Steps to Launch the Meta-analysis Job

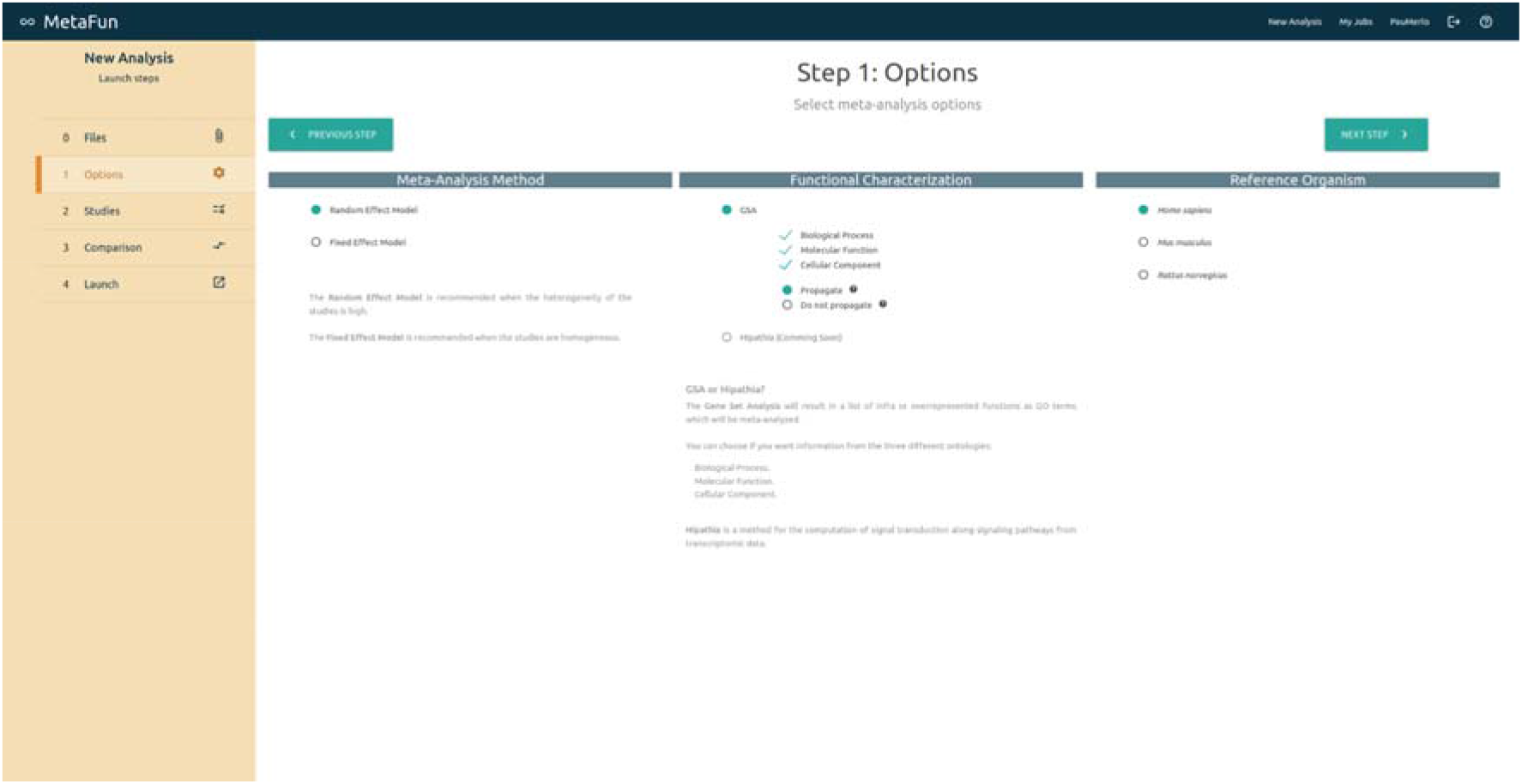
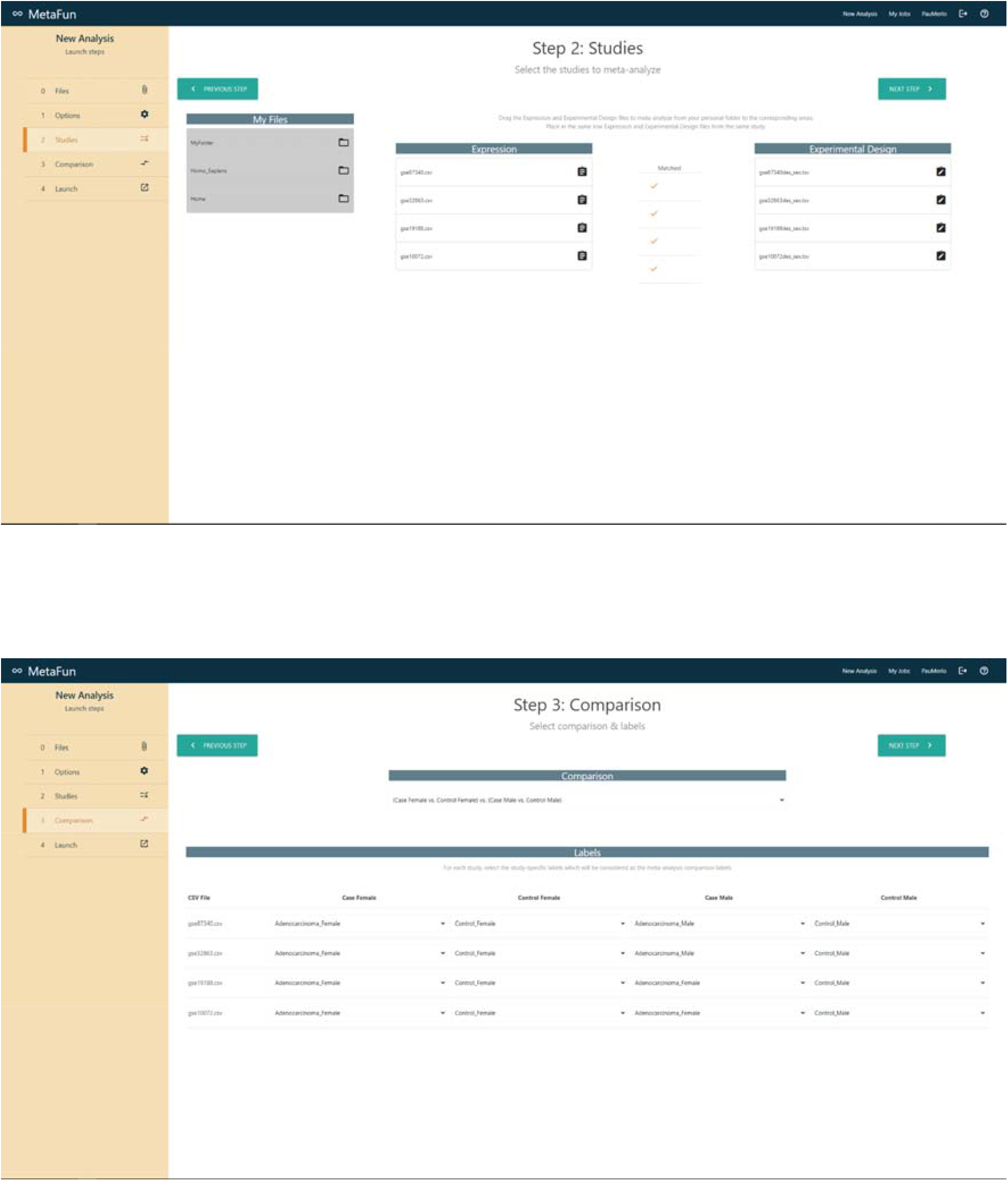
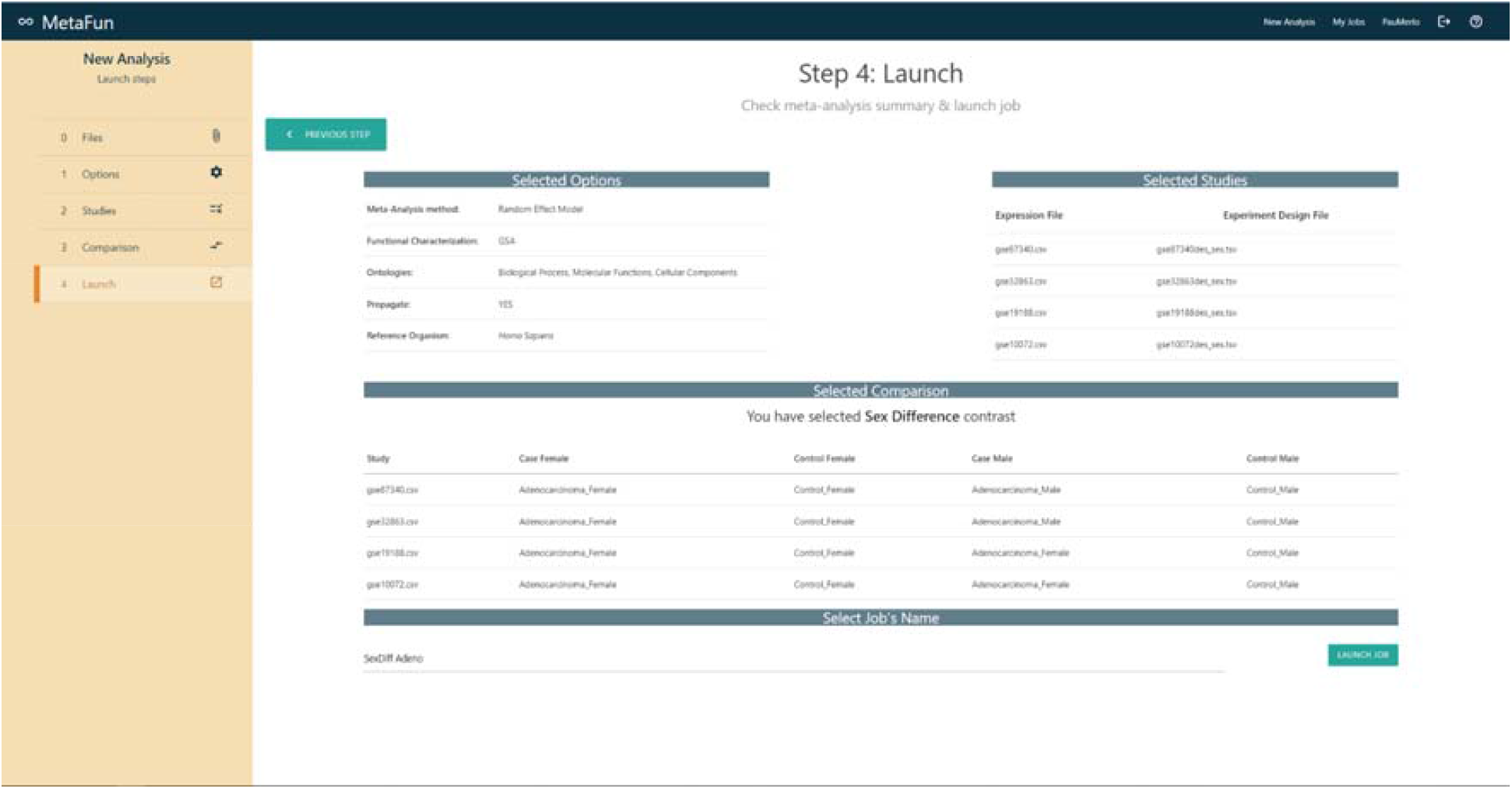

## Results

Below, we display the results generated by MetaFun in this use case for each of the sections described above (1. Analysis Summary, 2. Exploratory Analysis, 3. Differential Expression, 4. Gene Set Analysis, 5. Meta-analysis),

### 1. Analysis Summary

A summary of the results at the distinct stages of the bioinformatics analysis strategy:

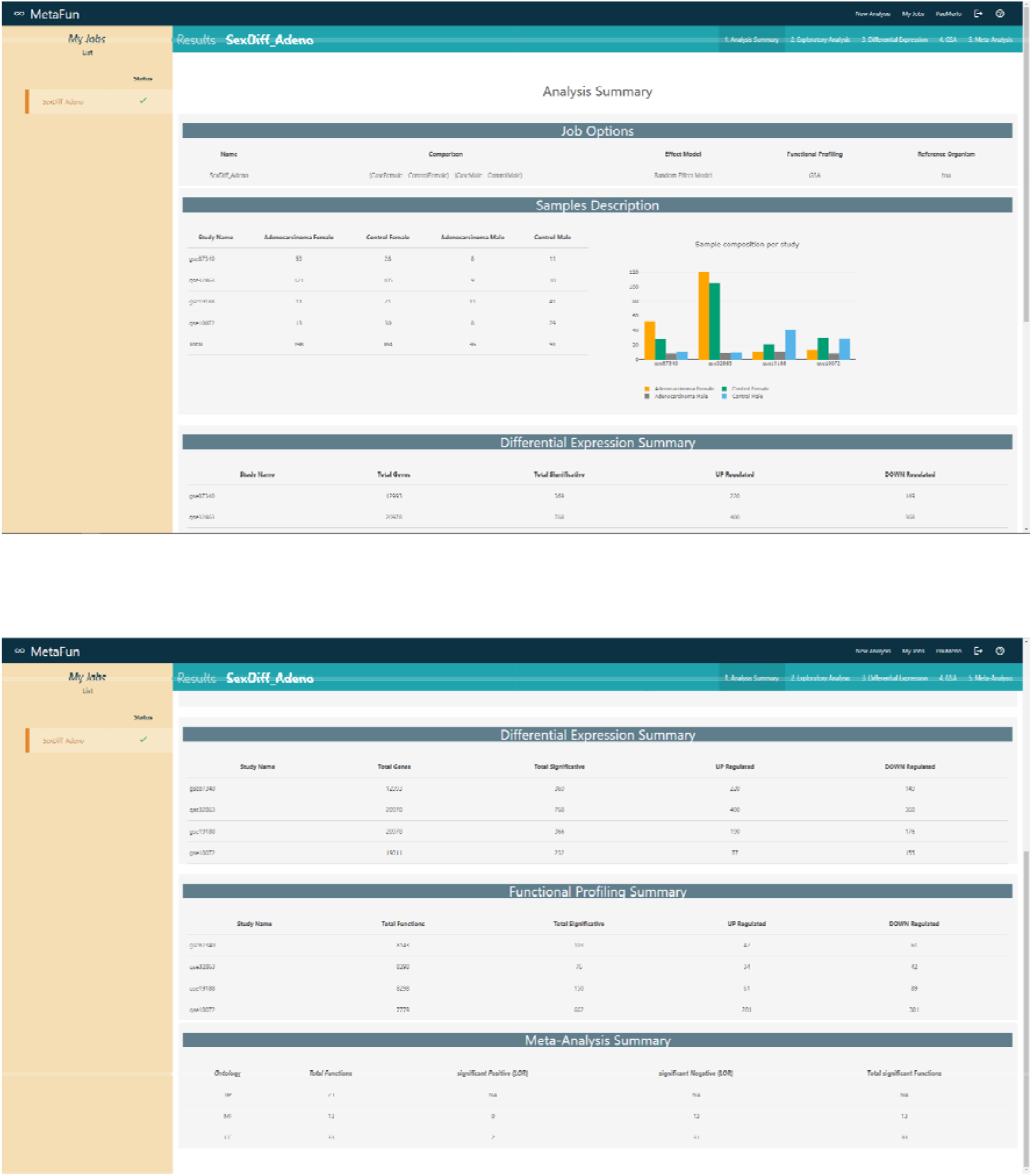

### 2. Exploratory Analysis

PCA, clustering, and boxplots are used to explore the expression levels of each of the samples in the selected studies:

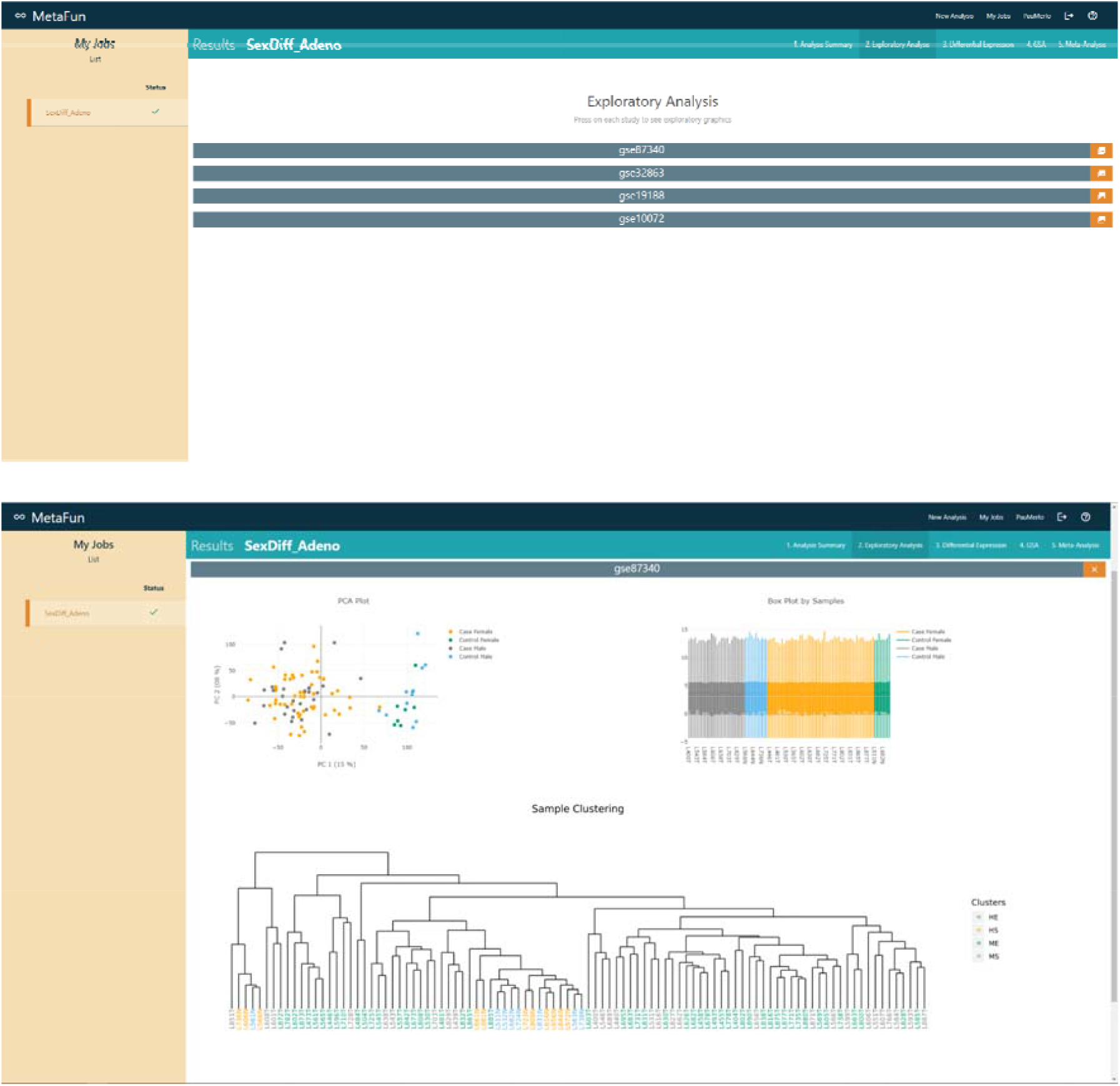

### 3. Differential Expression

The identification of genes showing differential expression by sex in affected patients for each study (Clicking on each link to the gene identifiers expands their biological information):

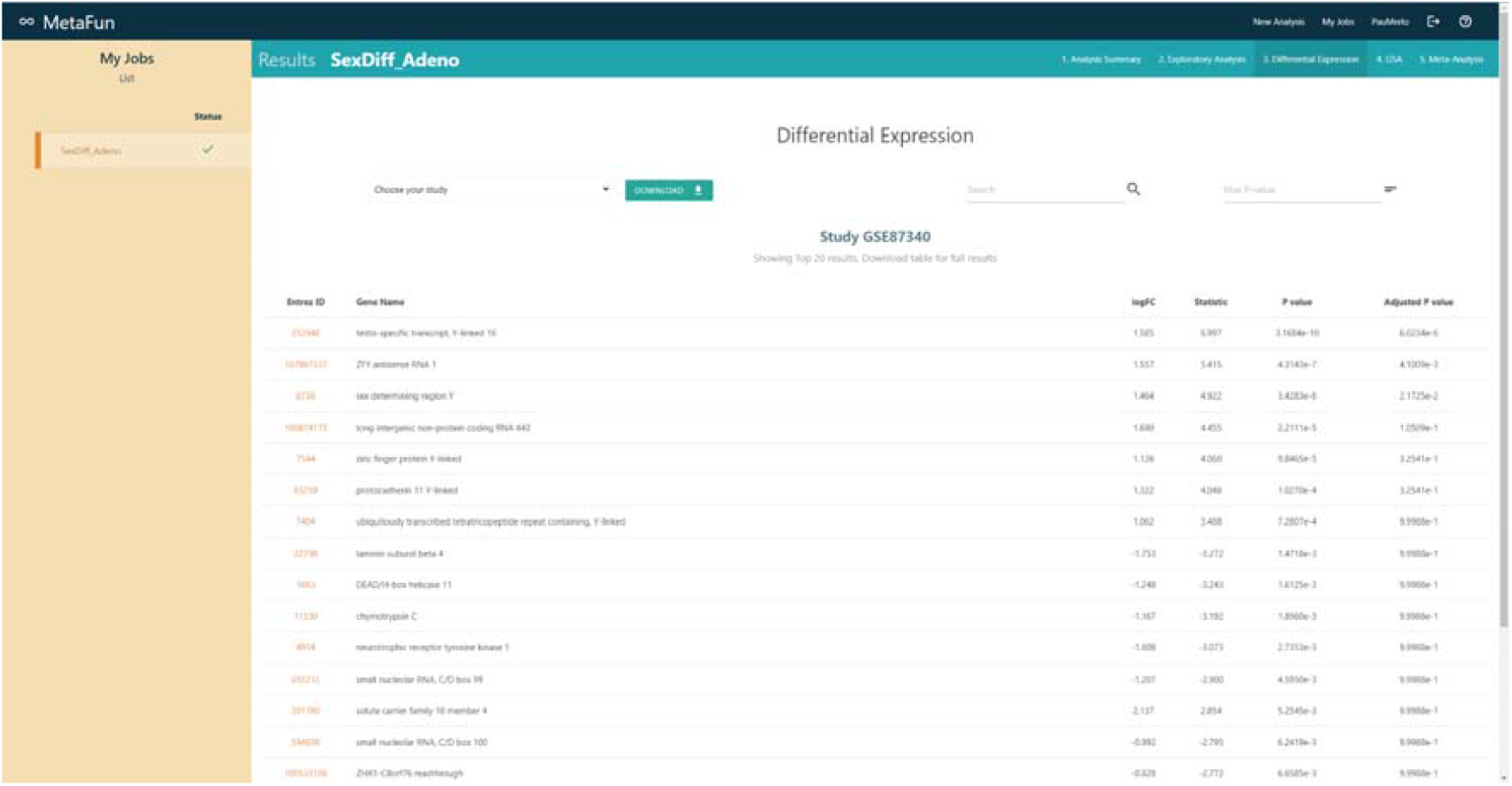

### 4. Gene Set Analysis (GSA)

Functional characterization of the differential expression results identifies those functions more active in males and females (Clicking on each link to the identifiers expands the information for each significant function):

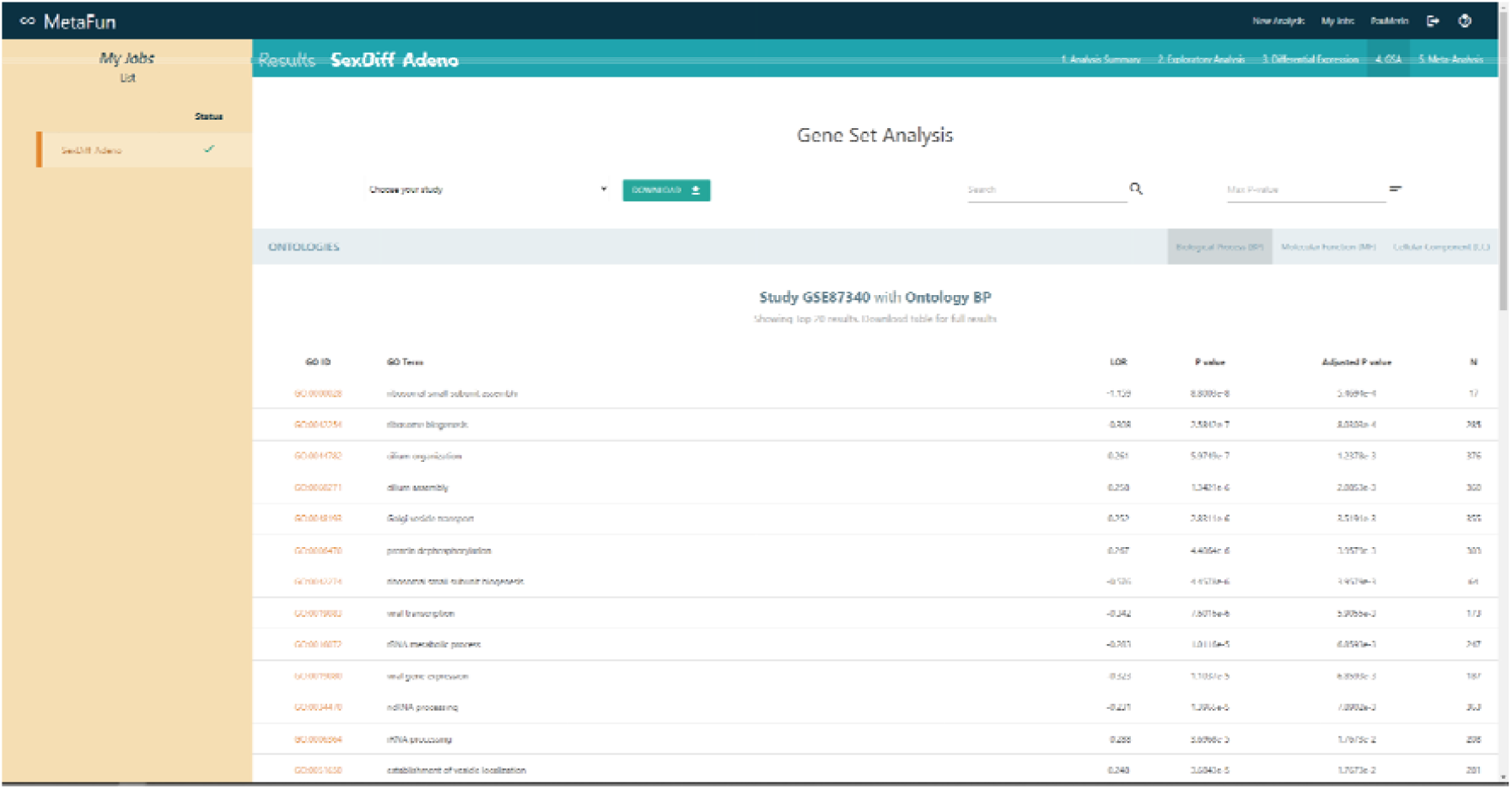

### 5. Meta-analysis

Demonstration of MetaFun functions and the pathways activated in evaluated studies (Clicking on the information icon leads to detailed information on these significant functions):

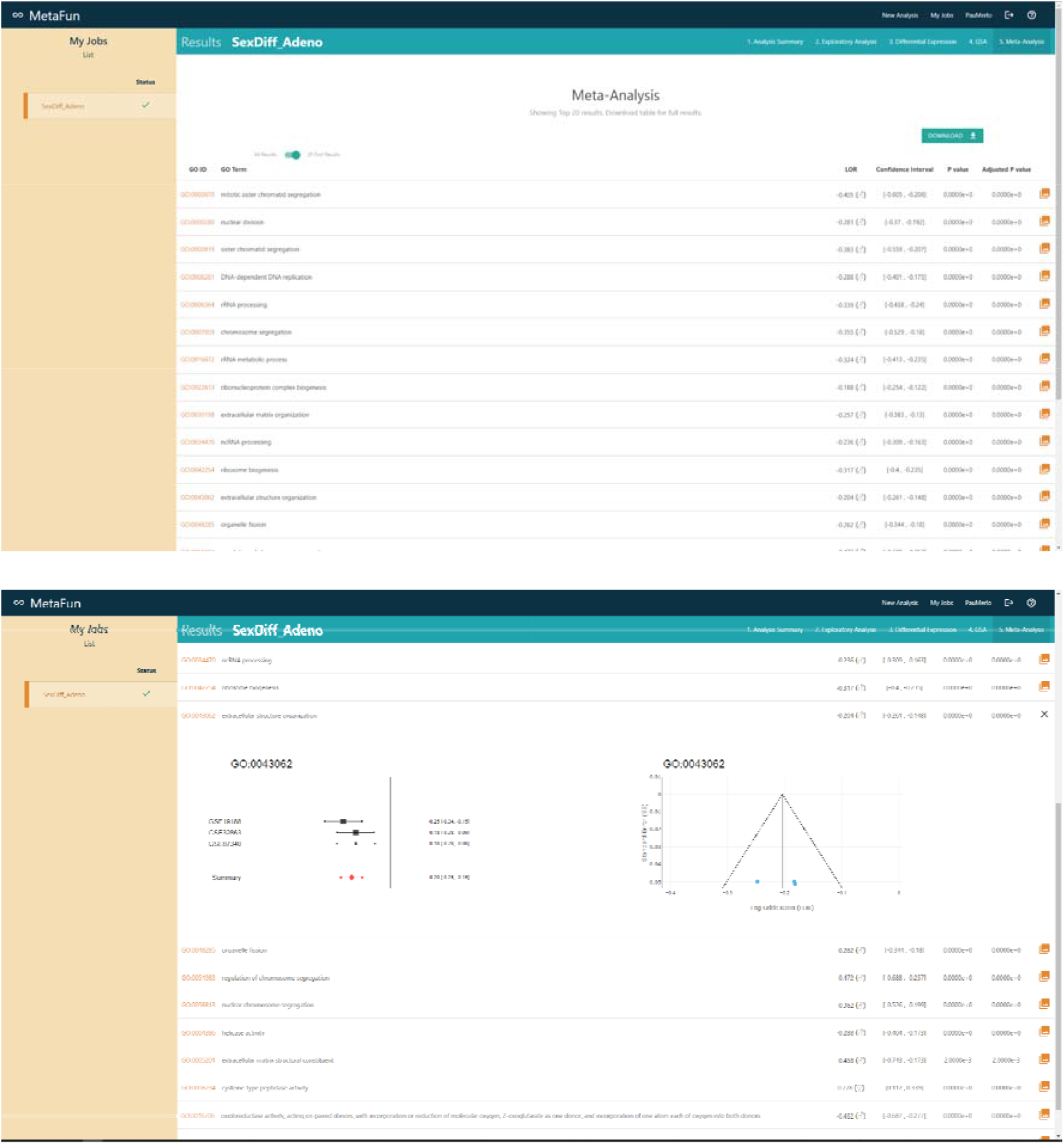

## References

1. Ballestri S. et al. NAFLD as a Sexual Dimorphic Disease: Role of Gender and Reproductive Status in the Development and Progression of Nonalcoholic Fatty Liver Disease and Inherent Cardiovascular Risk. Advances in Therapy. 2017; 34: 1291–1326. URL 10.1007/s12325-017-0556-1

2. Yuan Y, Liu L, Chen H, et al. Comprehensive Characterization of Molecular Differences in Cancer between Male and Female Patients. Cancer cell. 2016; 29(5): 711–722. doi:10.1016/j.ccell.2016.04.001

3. Barrett T, Wilhite SE, Ledoux P, Evangelista C, Kim IF, Tomashevsky M, Marshall KA, Phillippy KH, Sherman PM, Holko M, Yefanov A, Lee H, Zhang N, Robertson CL, Serova N, Davis S, Soboleva A. NCBI GEO: archive for functional genomics data sets--update. Nucleic Acids Res. 2013 Jan;41(Database issue):D991–5

4. Grossman RL, Heath AP, Ferretti V, Varmus HE, Lowy DR, Kibbe WA, Staudt LM. Toward a Shared Vision for Cancer Genomic Data. New England Journal of Medicine. 2016; 375:12, 1109-1112. doi: 10.1056/NEJMp1607591

5. Casanova Ferrer F, Pascual M, Hidalgo MR, Malmierca-Merlo P, Guerri C, García-García F. Unveiling Sex-Based Differences in the Effects of Alcohol Abuse: A Comprehensive Functional Meta-Analysis of Transcriptomic Studies. Genes. 2020; 11(9):1106. 10.3390/genes11091106

6. Pérez-Díez I, Hidalgo MR, Malmierca-Merlo P, Andreu Z, Romera-Giner S, Farràs R, de la Iglesia-Vayá M, Provencio M, Romero A, García-García F. Functional Signatures in Non-Small-Cell Lung Cancer: A Systematic Review and Meta-Analysis of Sex-Based Differences in Transcriptomic Studies. Cancers. 2021; 13(1):143. 10.3390/cancers13010143

7. Català-Senent JF, Hidalgo MR, Berenguer M, Parthasarathy G, Malhi H, Malmierca-Merlo P, de la Iglesia-Vayá M, García-García F. Hepatic Steatosis and Steatohepatitis: A Functional Meta-analysis of Sex-based Differences in Transcriptomic Studies. Biology of Sex Differences 2021, 12:29; 10.1186/s13293-021-00368-1

8. Plotly Technologies Inc. Collaborative data science. Montréal, QC, 2015. https://plot.ly

9. Ritchie ME, Phipson B, Wu D, Hu Y, Law CW, Shi W, Smyth GK. limma powers differential expression analyses for RNA-sequencing and microarray studies. Nucleic Acids Research. 2015; Volume 43, Issue 7, 20, Page e47,10.1093/nar/gkv007

10. Subramanian A, Tamayo P, Mootha VK, Mukherjee S, Ebert BL, Gillette MA, Paulovich A, Pomeroy SC, Golub TR, Lander ES, Mesirov JP. Gene set enrichment analysis: A knowledge-based approach for interpreting genome-wide expression profiles, Proceedings of the National Academy of Sciences Oct 2005, 102 (43) 15545–15550; doi: 10.1073/pnas.0506580102

11. Ashburner M, Ball CA, Blake JA, et al. Gene ontology: tool for the unification of biology. The Gene Ontology Consortium. Nat Genet. 2000;25(1):25–29. doi:10.1038/75556

12. Montaner D, Dopazo J. Multidimensional Gene Set Analysis of Genomic Data. PLoS One. 2010; 5(4), e10348. doi: 10.1371/journal.pone.0010348

13. Sayers, E. W., Bolton, E. E., Brister, J. R., Canese, K., Chan, J., Comeau, D. C., Connor, R., Funk, K., Kelly, C., Kim, S., Madej, T., Marchler-Bauer, A., Lanczycki, C., Lathrop, S., Lu, Z., Thibaud-Nissen, F., Murphy, T., Phan, L., Skripchenko, Y., Tse, T., ⃛ Sherry, S. T. (2022). Database resources of the national center for biotechnology information. Nucleic acids research, 50(D1), D20–D26. 10.1093/nar/gkab1112

14. Binns, D., Dimmer, E., Huntley, R., Barrell, D., O’Donovan, C., & Apweiler, R. (2009). QuickGO: a web-based tool for Gene Ontology searching. Bioinformatics (Oxford, England), 25(22), 3045–3046. 10.1093/bioinformatics/btp536

15. Viechtbauer W. Conducting meta-analyses in R with the metafor package. Journal of Statistical Software. 2010; 36(3), 1–48. https://www.jstatsoft.org/v36/i03/

16. https://www.mongodb.com/

17. Jain N, Mangal P & Mehta D. AngularJS: A modern MVC framework in JavaScript. Journal of Global Research in Computer Science. 2014; 5(12), pp.17–23

18. Wickham H. ggplot2: Elegant Graphics for Data Analysis. Springer-Verlag New York. 2016. ISBN 978-3-319-24277-4, https://ggplot2.tidyverse.org

19. Holm, S. (1979). A Simple Sequentially Rejective Multiple Test Procedure. Scandinavian Journal of Statistics, 6(2), 65–70. http://www.jstor.org/stable/4615733

20. Pérez-Díez I, Hidalgo MR, Malmierca-Merlo P, Andreu Z, Romera-Giner S, Farràs R, de la Iglesia-Vayá M, Provencio M, Romero A, García-García F. (2021). Functional Signatures in Non-Small-Cell Lung Cancer: A Systematic Review and Meta-Analysis of Sex-Based Differences in Transcriptomic Studies. Cancers (Basel). 2021 Jan 5;13(1):143.

